# Prediction of a plant intracellular metabolite content class using image-based deep learning

**DOI:** 10.1101/488783

**Authors:** Neeraja M Krishnan, Binay Panda

## Abstract

Plant-derived secondary metabolites play a vital role in the food, pharmaceutical, agrochemical and cosmetic industry. Metabolite concentrations are measured after extraction, biochemistry and analyses, requiring time, access to expensive equipment, reagents and specialized skills. Additionally, metabolite concentration often varies widely among plants, even within a small area. A quick method to estimate the metabolite concentration class (high or low) will significantly help in selecting trees yielding high metabolites for the metabolite production process. Here, we demonstrate a deep learning approach to estimate the concentration class of an intracellular metabolite, azadirachtin, using models built with images of leaves and fruits collected from randomly selected *Azadirachta indica* (neem) trees in an area spanning >500,000 sqkms and their corresponding biochemically measured metabolite concentrations. We divided the input data randomly into training- and test-sets ten times to avoid sampling bias and to optimize the model parameters during cross-validation. The training-set contained >83,000 fruit and >86,000 leaf images. The best models yielded prediction errors of 19.13% and 15.11% (for fruit), and 8% and 26.67% (for leaf), each, for low and high metabolite classes, respectively. We further validated the fruit model using independently collected fruit images from different locations spanning nearly 130,000 sqkms, with 70% accuracy. We developed a desktop application to scan offline image(s) and a mobile application for real-time utility to predict the metabolite content class. Our work demonstrates the use of a deep learning method to estimate the concentration class of an intracellular metabolite using images, and has broad applications and utility.

## Background

Measuring the concentration of an analyte, such as metabolite, enzyme, protein or any other chemical moiety within plants, animals and microbial cells is a frequent practice in biology. To do this, chemical, biochemical, immunological or imaging-based methods are usually followed, where each method provides a different type of readout. Although accurate and precise, the currently employed analytical methods require extensive sample handling and preparation time, expensive reagents and equipment, and specialized skills. In circumstances where knowing the exact concentration of the intracellular metabolite is not necessary, and a quick and rough estimate of its concentration class (either high or low) is enough, use of conventional measurement methods although unnecessary is currently unavoidable. A method that provides a quick readout of the analyte concentration class, however ideal in such cases, is currently not available.

Metabolites, primary and secondary, are intermediate products of metabolic reactions catalyzed by enzymes. Examples of some primary metabolites are amino acids, vitamins, organic acids, and nucleotides. Living cells synthesize most of the primary metabolites essential for their growth. Secondary metabolites, especially those derived from microbial and plant sources, such as drugs, fragrances, dyes, pigments, pesticides and food additives, are not required for the cell’s primary metabolic processes but have a wide range of application in agriculture and pharmaceutical industry. The commercial importance of secondary metabolites has invoked the exploration of producing bioactive plant metabolites synthetically in the laboratory, either by total synthesis, plant tissue culture or metabolic engineering [1].

Machine learning and artificial intelligence-based methods have recently found their utility in biology [2, 3]. Algorithms based on deep learning can predict patterns from raw biological data (genomic data, images, proteins etc.) and help to generate biological inferences. Deep learning methods outperform conventional methods of image classification, such as principal components identification, and their usage in Support Vector Machines and Random Forest classifiers [4]. They are, therefore, more suitable for image-based analyses.

In the current manuscript, we focussed on a plant-based commercially important secondary metabolite, azadirachtin, a potent and versatile tetranotriterpenoid found predominantly in the fruits, and to a lesser degree in the leaves, of the tree, *Azadirachta indica* (neem) [5]. We used an image-based deep learning approach utilizing convolutional neural networks (CNNs) to learn from the original images of leaves and fruits from the sampled trees and the image distortions, with a total of nearly 170,000 images, alongside the biochemically measured concentrations of azadirachtin, to build models that can predict the metabolite concentration class (high or low) for an incoming new leaf or fruit.

## Methods

The deep learning approach described here used CNNs to learn from >83,000 fruit and >86,000 leaf images, and the azadirachtin content classes derived from biochemically measured azadirachtin content, to build predictive fruit and leaf models. The models were optimized based on their performance in blind test sets. The flow diagram in Figure 1 describes the training set and training criteria used in building models, and the different thresholds for azadirachtin class (azaL – low azadirachtin and azaH – high azadirachtin) prediction accuracies used in short-listing and analyzing models.

**Figure 1.**
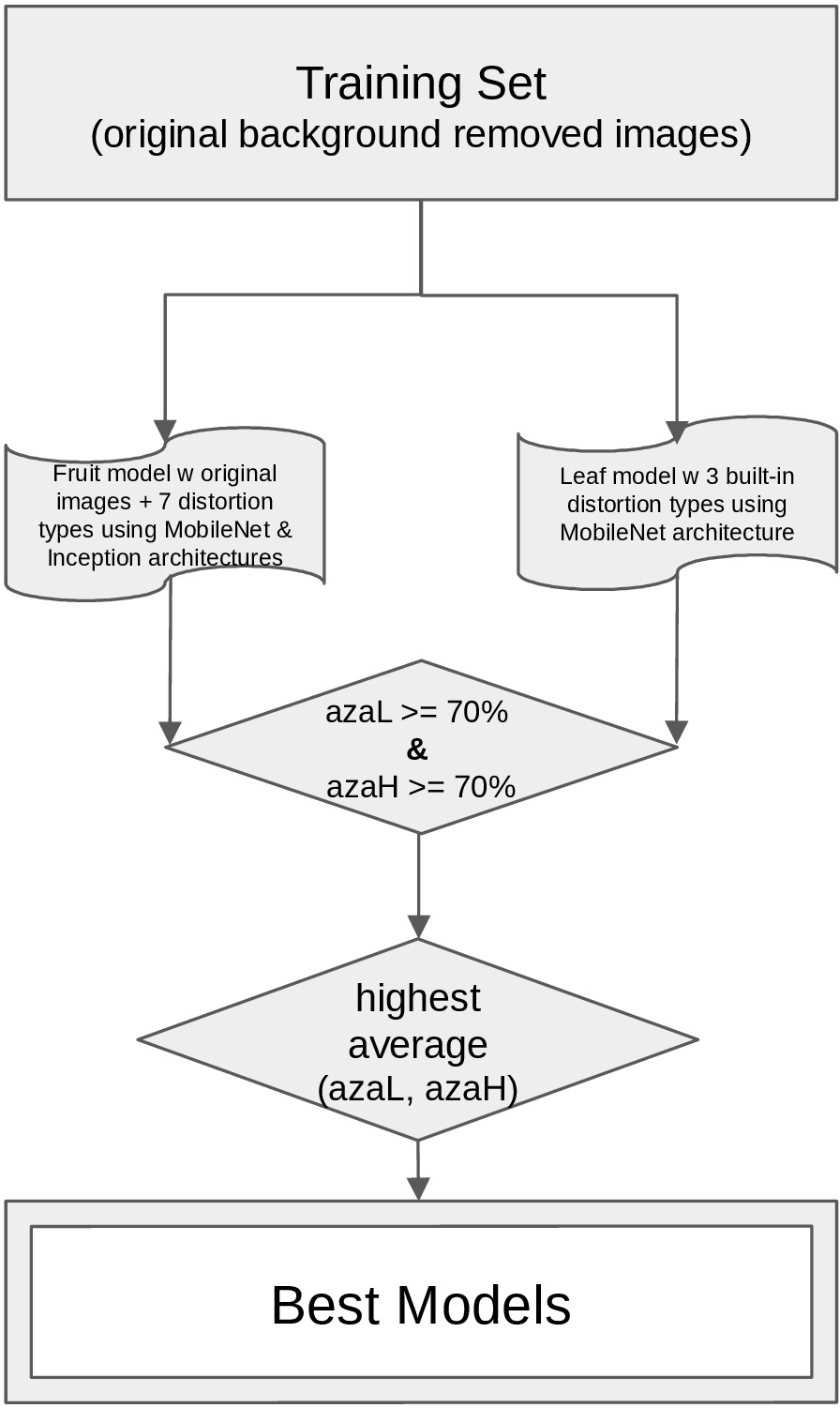
Flow diagram describing the training set and training criteria used in building models. Different thresholds used for azadirachtin class (azaL – low azadirachtin and azaH – high azadirachtin) prediction accuracies in short-listing and finalizing models are also provided.

### Plant material collection and Metabolite measurement

We randomly sampled trees in an area covering >500,000 sq kms and collected five leaves and five fruits per tree. Both leaves and fruits were photographed and Azadirachtin (azadirachtin A and azadirachtin B, together) was measured using reverse-phase columns on High Performance Liquid Chromatography (HPLC) instrument with azadirachtin (A+B) analytical standards (EID Parry, India).

### Image analyses

In order to remove bias originating from the background, the original images were subjected to background removal using an online editing tool, Background Burner, personal edition (https://burner.bonanza.com/). This step was performed to base the classification on the foreground, and not the variable background. Training with background-removed images has been shown previously to yield high prediction accuracy [6]. For real-time prediction purpose, the incoming image background removal option was incorporated using an open source Android image background cutting library called Android-CutOut v0.1.2 (https://github.com/GabrielBB/Android-CutOut) within the mobile app, prior to classification. This was done using two options: automatically by touching parts of the background, using the ‘magic wand’ option, and manually by dragging the finger around the screen using the ‘pencil’ and ‘zoom’ options. Depending on the contrast between the background and foreground of the image, automatic only or both automatic and manual options can be used. For example, for a leaf image captured against a blue background, the ‘magic wand’ tool can remove the blue background, in one step. However, for a leaf image captured against a background, which has the leaf ‘s green color in it, touching the green color in the background with the magic wand tool, will remove it from the foreground as well. This is where the manual option comes in handy to selectively remove the green parts of the residual background. We used this library in the mobile app because it was compatible and easy-to-integrate with the TensorFlow based classification workflow. The images were then used for azadirachtin classification. In the desktop version, we processed the incoming images with Background Burner before using AZA classifier.

In addition to the original images, we introduced several distortions to make a robust model for training. We did this by using a Python module, Augmentor [7], to augment the images with 7 types of distortions: Rotate, Zoom, Flip, Blur, Crop, Brightness and Resize. The options used with the Augmentor module to create these distortions are summarised in Table 1. Representative images of leaves and fruits, before and after distortions are shown in Figure 2.

**Table 1.**
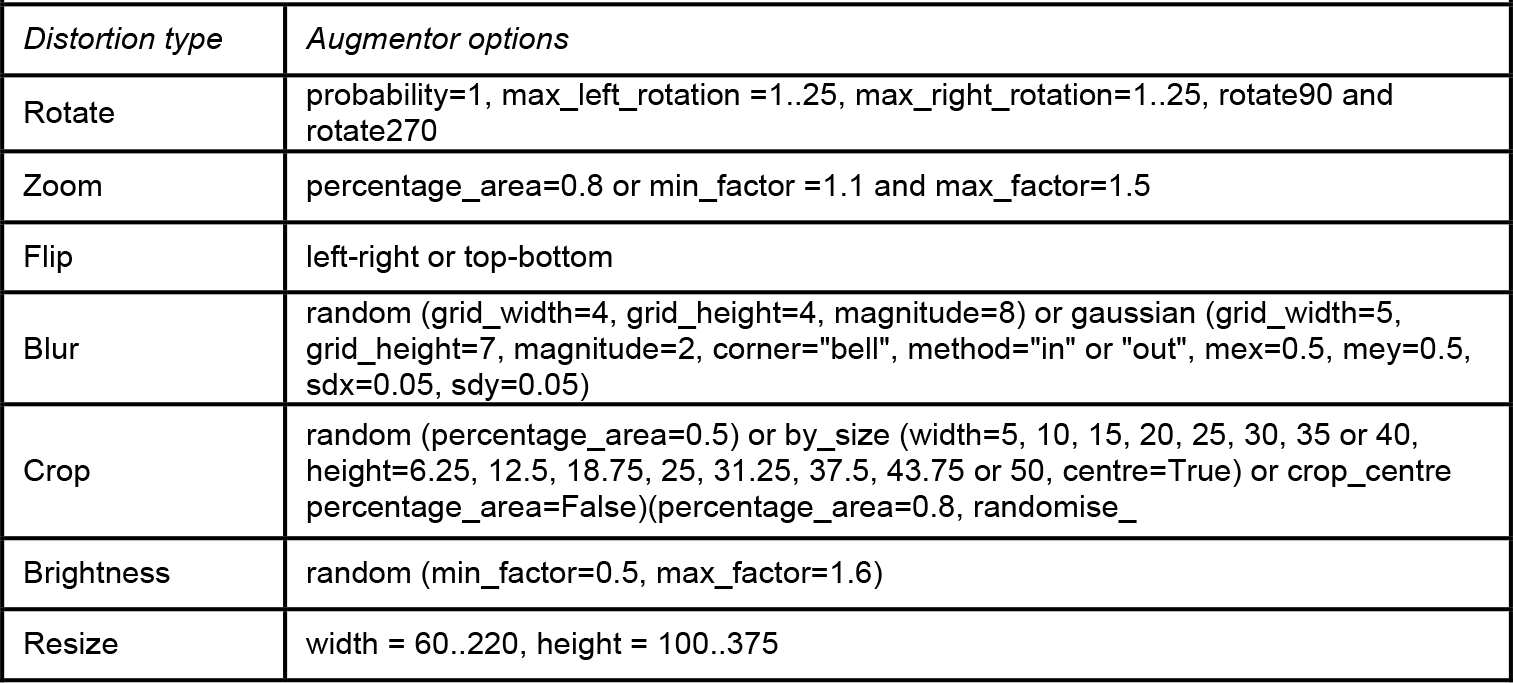
Options used for making the image distortions.

**Figure 2.**
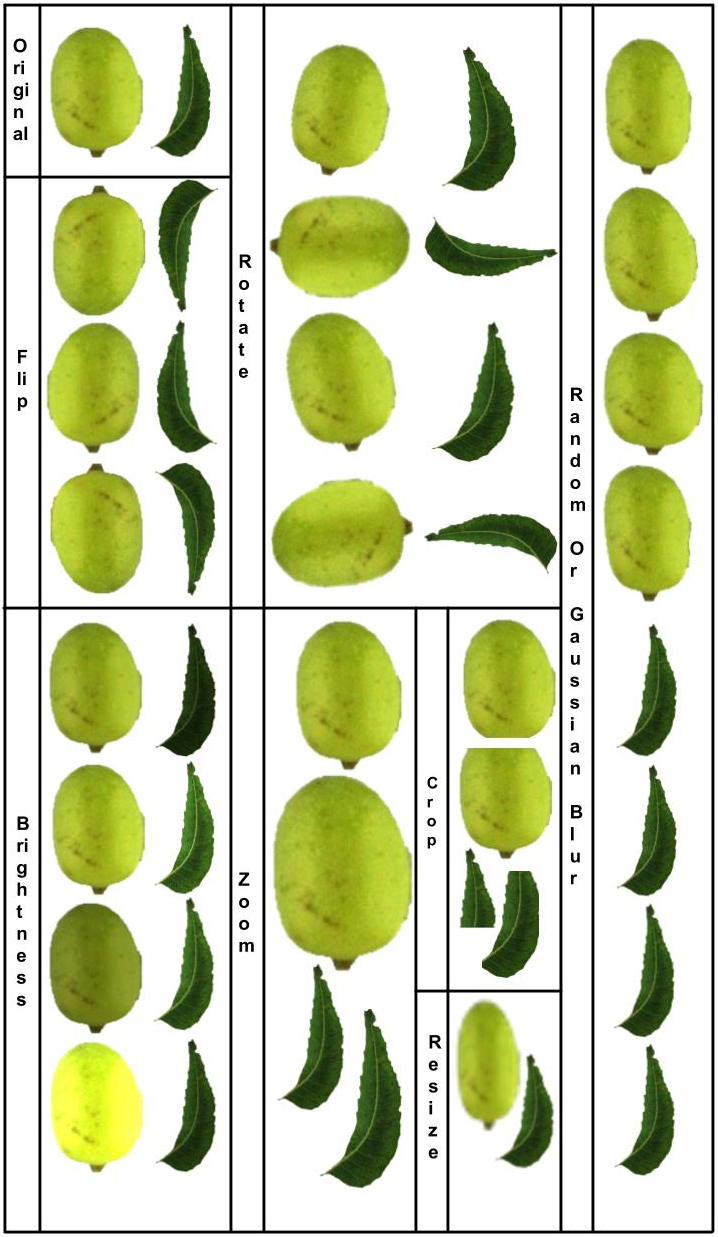
Representative image of a leaf and a fruit subjected to seven synthetic distortions. Depending on the type of distortion, variable numbers of images were created post distortion. Multiple representative images are shown for some distortions in the figure to display the range of effects.

### Learning and Test Set Classification

As azadirachtin derived from seed kernel is considered to be the gold standard, we used seed kernel-derived metabolite values to set concentration class as either “low (azaL)” or “high (azaH)”. Once the minimum and the maximum azadirachtin values were determined, the average (avg) was calculated as the mid-point of those min-max extremes. An azadirachtin class was set as ‘low’ and ‘high’ when the measured concentrations were below avg − 0.1avg and above avg + 0.1avg, respectively. Accordingly, an azadirachtin value above 0.65% was determined as ‘high’, and a value below 0.53% was determined as ‘low’. The sample numbers for the azaL and azaH classes in ten learning and test sets randomly generated for 10-fold cross validation are summarised in Table 2. The corresponding images post distortions were also divided into learning and test sets for training and classification. For each learning set, the azaL and azaH classification sensitivity values were calculated as the average value across the ten test sets.

**Table 2.** Division of fruit/leaf images into ten random pairs of learning and test sets for the azaL and azaH classes.

| Table 2 |  |  |  |  |  |
| --- | --- | --- | --- | --- | --- |
| *azaL (< 0.53 %)* |  |  | *azaH (> 0.65 %)* |  |  |
| **Set** | **Set ID** | **Number of images** | **Set** | **Set ID** | **Number of images** |
| Learning | 1 | 445 | Learning | 1 | 295 |
|  | 2 | 445 |  | 2 | 290 |
|  | 3 | 445 |  | 3 | 295 |
|  | 4 | 445 |  | 4 | 290 |
|  | 5 | 445 |  | 5 | 295 |
|  | 6 | 445 |  | 6 | 290 |
|  | 7 | 445 |  | 7 | 295 |
|  | 8 | 445 |  | 8 | 290 |
|  | 9 | 445 |  | 9 | 295 |
|  | 10 | 450 |  | 10 | 290 |
| Test | 1 | 50 | Test | 1 | 30 |
|  | 2 | 50 |  | 2 | 35 |
|  | 3 | 50 |  | 3 | 30 |
|  | 4 | 50 |  | 4 | 35 |
|  | 5 | 50 |  | 5 | 30 |
|  | 6 | 50 |  | 6 | 35 |
|  | 7 | 50 |  | 7 | 30 |
|  | 8 | 50 |  | 8 | 35 |
|  | 9 | 50 |  | 9 | 30 |
|  | 10 | 45 |  | 10 | 35 |

### Algorithm and Framework

We named the tool ‘AZA classifier’ based on the test case, herein, to classify azadirachtin content class. AZA classifier (desktop version) is developed using JAVA swing GUI [8] for its native look and is backed up with Linux shell scripting and Python3 in its backend. The desktop version of AZA classifier was built to classify azadirachtin content class (low or high) for a single leaf or fruit image, or bulk process multiple images saved in a folder (Figure 3A). The results from the desktop app were bulk exported into an Excel sheet, while storing the image identity, with predicted azadirachtin class, and prediction confidence (probability ranging from 0 to 1) and analyzed further. We used TensorFlow 1.6.0 packages [9], an open-source framework for building deep learning neural networks [10], in Python3 to build predictive models that could classify images of leaves and fruits. We used the MobileNet architecture [11] that overcomes the challenges in visual recognition of images, particularly for on-mobile device or embedded applications to run the models effectively where resources like computation, power and space are limited. It is a family of mobile-first computer vision models for TensorFlow, designed specifically for an effective increase in accuracy, with limited resources for an on-device or embedded application. They are actually small, low-power, low-latency models designed to deal with limited resources in a number of use cases. The latest release (1.13) consists of model definition for MobileNets in TensorFlow using TF-Slim in addition to 16 pre-trained ImageNet classification checkpoints for use in mobile projects of any size. The Inception deep convolutional architecture, introduced as GoogLeNet, was the named as the Inception v1 architecture [12]. It was later improvised, first by batch normalization - Inception v2 [13], and then by additional factorization ideas - Inception v3 [14]. In addition to MobileNet, we also explored the Inception v3 architecture [14]. The mobile version of AZA classifier (Android app) was built using Android Studio 3.0 [15], compiled and run based on TensorFlow Android version 1.6.0. In addition to taking still leaf and fruit images, it is also capable of a live or real-time classification of the azadirachtin content class by pointing the camera of the mobile device to a leaf or a fruit while they are still attached to a tree (Figure 3B). After the user captures a leaf or a fruit image with the mobile camera, the background is removed before the image is used for the classification of azadirachtin. This function makes the mobile app capable of classifying real world images. A “Save” option in the app enables the user to capture an image along with the azadirachtin classification while providing an identifier to the captured image. In cases of fluctuation in the image capture process, the user has the provision to capture the same image multiple times until a satisfactory image quality is obtained. The effects of different zoom levels, different angles, and different light conditions have already been explored and factored in during the training process while considering augmentations of the images.

**Figure 3.**
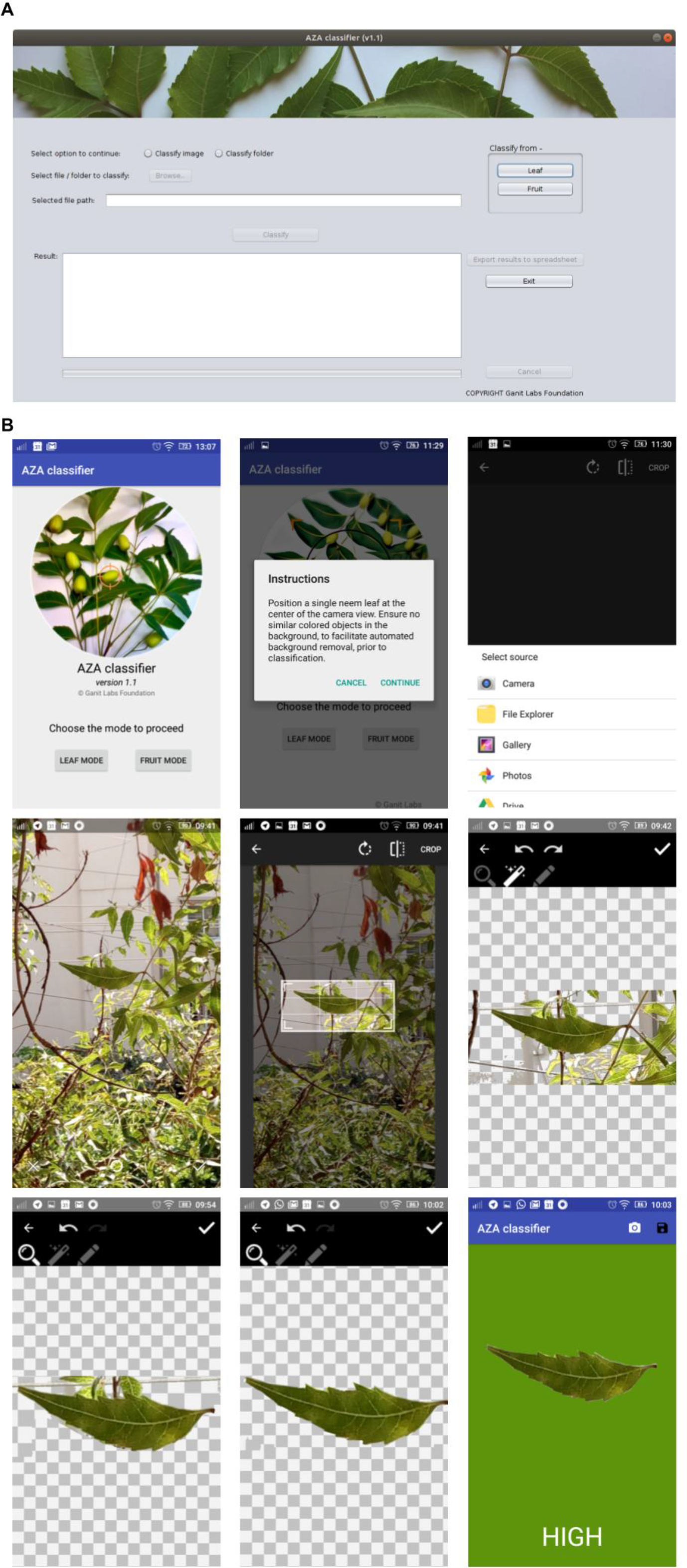
Interface of the AZA classifier app in the A. Desktop and B. Mobile versions. In the mobile app, the crop and background removal steps with a real-world image (still attached to a tree) are demonstrated (B).

### Training set and model testing using TensorFlow library with CNN architecture

Using TensorFlow, we built azadirachtin prediction models for 17 types of configurations including 16 MobileNet (pre-trained deep CNN network versions for width multipliers of 0.25, 0.50, 0.75 and 1.0, and image input sizes of 128, 160, 192, and 224) and Inception (pre-trained deep CNN network) v3 [14] models, using pre-trained weights. Each model was trained for 500, 1000, 1500, 2000, 2500, 3000, 3500, 4000, 4500 and 5000 training steps.

Images were divided into azaL and azaH classes. The models were trained and tested with all image augmentations. In addition, the models were also trained and tested for each augmentation individually, and without any augmentation, for comparison.

The classification sensitivity at which there was at least one model for both azaL and azaH classes was estimated for each individual distortion type. Following this, 160 MobileNet and 10 Inception models (with the ten training steps mentioned previously) were constructed after training with all the 8 types of images pooled together, and tested with images not used during training. A learning rate of 0.01 was used for training with cross-entropy as the loss function. For the leaf, the images, post background removal, were also distorted internally (Crop - 30%, Scale - 30% and Flip - left and right) using the python module ‘retrain’ of TensorFlow library https://github.com/tensorow/hub/blob/master/examples/image_retraining/retrain.py, followed by training and classification using 160 MobileNet models. The fruit models from MobileNet and Inception architectures, with at least 80% classification sensitivities for both azaL and azaH classes were shortlisted. Likewise, the leaf models were shortlisted from TensorFlow’s inbuilt distortions while retraining using MobileNet, with at least 70% classification sensitivities for both classes. These thresholds were set as the highest common classification sensitivities for azaL and azaH classes. The final fruit and leaf models were chosen with the highest average classification sensitivity between azaL and azaH classes.

### Independent validation of the model

Images of randomly sampled fruits from independent locations covering nearly 130,000 sq kms, which were not used for training or testing, were used as a validation set.

## Results

### Images for the training set

We used seven distortions on the images post background removal (*see* Methods). The number of images used in training and testing for each distortion type are listed in Table 3. The image numbers were highest for the Rotation distortion type; augmented 60-fold over the original followed by 22-fold for the Crop distortion and 17-fold for the Resize distortion type. The fold-differences were correlated with the number of distortion possibilities and outcomes for each type. The total numbers of images used for training were 83,898 (48285 for azaL & 35613 for azaH) for fruits and 86,311 (51,495 for azaL & 34,816 for azaH) for leaves (Table 3).

**Table 3.** Number of fruit and leaf images used for the training and test sets after the backgrounds have been removed, and after individual distortions are applied. Total numbers of images used for training for classification (pooled) were 83,898 (48285 for azaL & 35613 for azaH) for fruits and 86,311 (51,495 for azaL & 34,816 for azaH).

| Table 3 |  |  |  |  |  |
| --- | --- | --- | --- | --- | --- |
| Distortion type | Organ | Training set |  | Test set |  |
|  |  | azaL | azaH | azaL | azaH |
| Background removal | fruit | 435 | 297 | 45 | 30 |
|  | leaf | 429 | 293 | 50 | 30 |
| Crop | fruit | 9570 | 6556 | 990 | 660 |
|  | leaf | 9438 | 6446 | 1100 | 660 |
| Brightness | fruit | 870 | 576 | 90 | 60 |
|  | leaf | 858 | 586 | 100 | 60 |
| Gaussian/random blur | fruit | 2610 | 1788 | 330 | 120 |
|  | leaf | 2574 | 2334 | 300 | 180 |
| Rotate | fruit | 25230 | 16038 | 2430 | 1620 |
|  | leaf | 27032 | 15823 | 2700 | 4320 |
| Flip | fruit | 1740 | 3802 | 180 | 300 |
|  | leaf | 1726 | 2888 | 200 | 320 |
| Zoom | fruit | 870 | 1192 | 180 | 120 |
|  | leaf | 1716 | 1172 | 200 | 120 |
| Resize | fruit | 6960 | 5364 | 810 | 540 |
|  | leaf | 7722 | 5274 | 900 | 540 |
| Pooled | fruit | 48285 | 35613 | 5055 | 3450 |
|  | leaf | 51495 | 34816 | 5550 | 6230 |

### Suitable fruit and leaf models

A training set pooling all image augmentations was constructed to build models using MobileNet and Inception architectures. Models were deemed suitable if they predicted with at least 70% classification sensitivity for azaL and azaH content classes (*Tables S1-S2*). As per these criteria, 15 fruit models were deemed suitable with the MobileNet architecture, and 7 with the Inception architecture. No leaf model passed these criteria, with either architecture. Augmented with three inbuilt distortions as a training set, six leaf models were deemed suitable, within the original background-removed test set of leaf images (*Table S3*). In comparison, for individual augmentations, there were 5 suitable fruit models for Brightness, 9 for Resize, 4 for Crop, 13 for Random/Gaussian blur, 18 for Rotate, 0 for Zoom and 28 for Flip, and 1 suitable leaf model for Brightness, 0 for Resize, Crop, Random/Gaussian blur, and Rotate, 5 for Zoom, and 2 for Flip distortion types, and for no augmentation, there were 15 and 4, suitable fruit and leaf models, respectively, with MobileNet architecture (*Table S4*).

### Final fruit and leaf models of metabolite classification

The thresholds for shortlisting fruit and leaf models trained with pooled augmented data, were individually set to be the highest common classification sensitivities for azaL and azaH classes, at 80% and 70%, respectively. Two fruit models and six leaf models passed these criteria. The final models were chosen from Inception v3 architecture, at 2500 training steps, for the fruit, with an average classification sensitivity of 82.88%, and from the MobileNet architecture, with built-in distortions while retraining, image resolution of 128, width multiplier of 0.50, at 2500 training steps, for the leaf, with an average classification sensitivity of 82.67% (Table 4). The leaf model derived fromatrainingsetaugmentedwiththree inbuilt distortion types (Crop, Flip and Scale) was less sensitive in classifying azadirachtin content class within a test set that accommodated four more distortion types, thus resulting in a slightly lower classification sensitivity of >70% for both azaL and azaH classes (Table 4). For Resize, Rotate and Crop distortion types, while the azaL classification sensitivity was 90.67%, 77.26% and 89.27%, respectively, the azaH classification sensitivity remained below 50% at 25.74%, 42.08% and 42.58%, respectively (Table 4).

**Table 4.** Final fruit (orange) and leaf (green) models.

| Table 4 |  |  |  |  |  |  |  |
| --- | --- | --- | --- | --- | --- | --- | --- |
| Image type | Architecture | Image resolution | Width multiplier | No. of training steps | azaL classification sensitivity (%) | azaH classification sensitivity (%) | Average classification sensitivity (%) |
| fruit | InceptionV3 | -na- | -na- | 500 | 82.81 | 81.56 | 82.18 |
| fruit | InceptionV3 | -na- | -na- | 2500 | 80.87 | 84.89 | 82.88 |
| leaf | MobileNet | 224 | 0.25 | 1000 | 70.00 | 80.00 | 75.00 |
| leaf | MobileNet | 224 | 0.25 | 4500 | 84.00 | 73.33 | 78.67 |
| leaf | MobileNet | 192 | 0.75 | 3000 | 76.00 | 70.00 | 73.00 |
| leaf | MobileNet | 192 | 1.00 | 5000 | 76.00 | 70.00 | 73.00 |
| leaf | MobileNet | 128 | 0.5 | 2500 | 92.00 | 73.33 | 82.67 |
| leaf | MobileNet | 128 | 0.75 | 500 | 86.00 | 73.33 | 79.67 |

### Using both models (fruit and leaf) for classification

The result for the metabolite class prediction always provides a binary (correct or not) output when a single model (either fruit or a leaf) is used. However, when both the models are used for classification, there are four possible prediction scenarios,: A. both the fruit and the leaf model classify the class correctly, B. only the fruit model classifies it correctly but the leaf model does not, C. only the leaf model classifies it correctly but the fruit model does not, and D. both the models fail to classify the class correctly, *i.e.*, both models misclassify. We visualized each of these prediction scenarios with our test set (Figure 4). We observed that 25 out of 40 azaH images are correctly classified by both the fruit and the leaf models, 10 were correctly classified by the fruit model alone but misclassified by the leaf model, 1 was misclassified by the fruit model but correctly classified by the leaf model, and 4 were misclassified by both the models (Figure 4). Out of the 64 azaL images, 52 were correctly classified by both fruit and leaf models, 4 were correctly classified by the fruit but not the leaf model and 8 were correctly classified by the leaf but not the fruit model. Table 5 enumerates the true positive and false positive numbers for these azaH and azaL images, while using the fruit model alone, or the leaf model alone, or combining the predictions from both the models.

**Figure 4.**
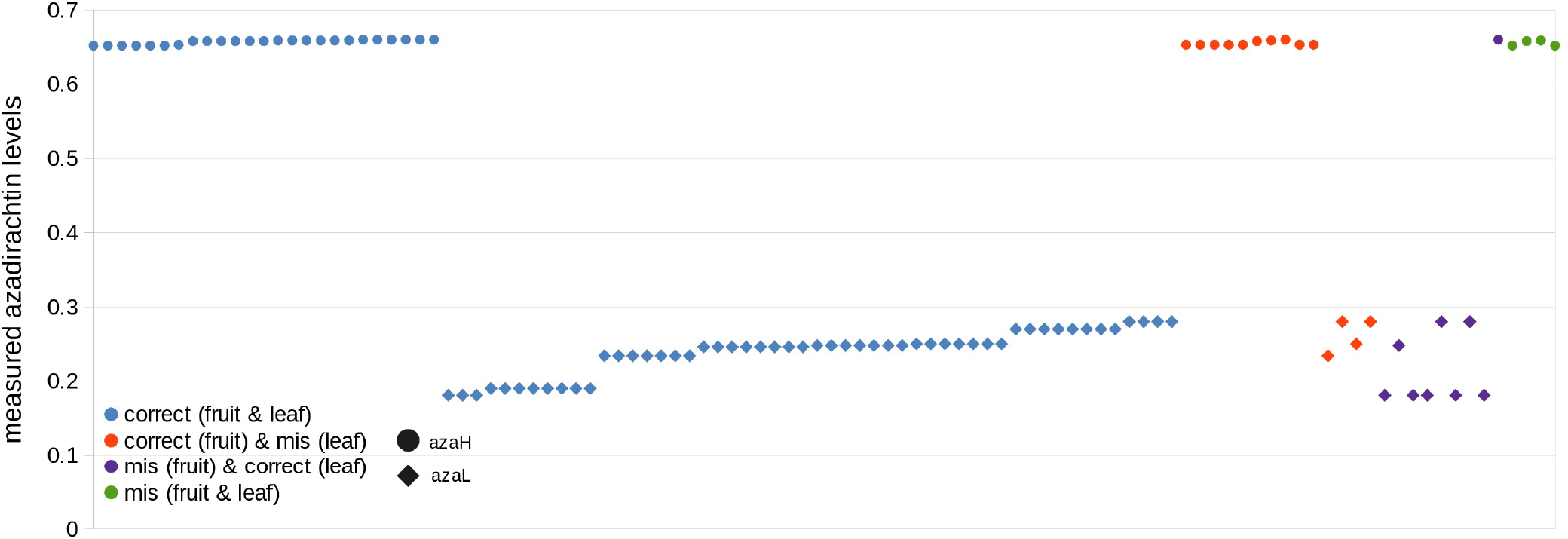
Azadirachtin classification using the fruit and the leaf models. Classification under all the four scenarios are visualized against measured azadirachtin levels: (1) where both the fruit and the leaf models correctly classify the metabolite class, (2) where the fruit model alone but not the leaf model classifies correctly, (3) where the leaf model alone but not the fruit model classifies correctly and finally, (4) where both the fruit and the leaf model mis (misclassify).

**Table 5.**
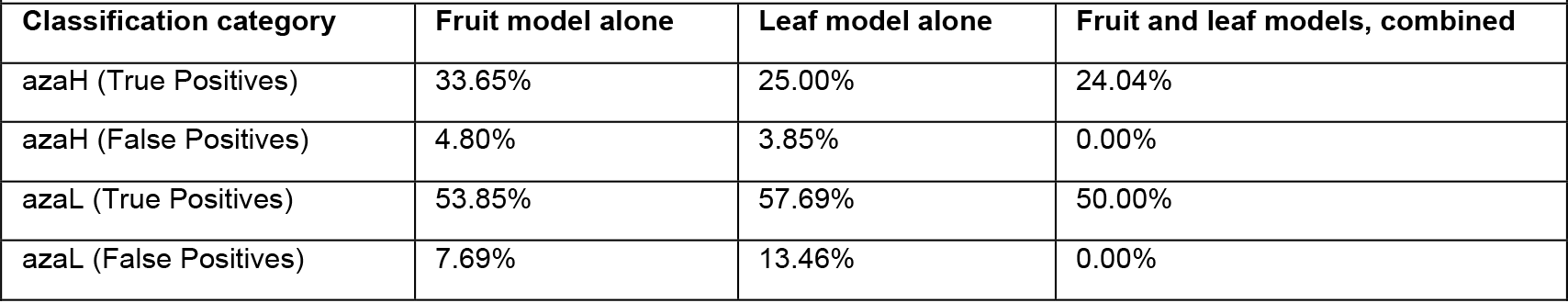
Percentage of true and false positives while classifying with the fruit model alone, the leaf model alone, and with both the fruit and the leaf models, combined.

### Independent validation of fruit models

The final fruit model was validated in an independent set of 115 fruit images, with 70% classification sensitivity (Table 6).

**Table 6.**
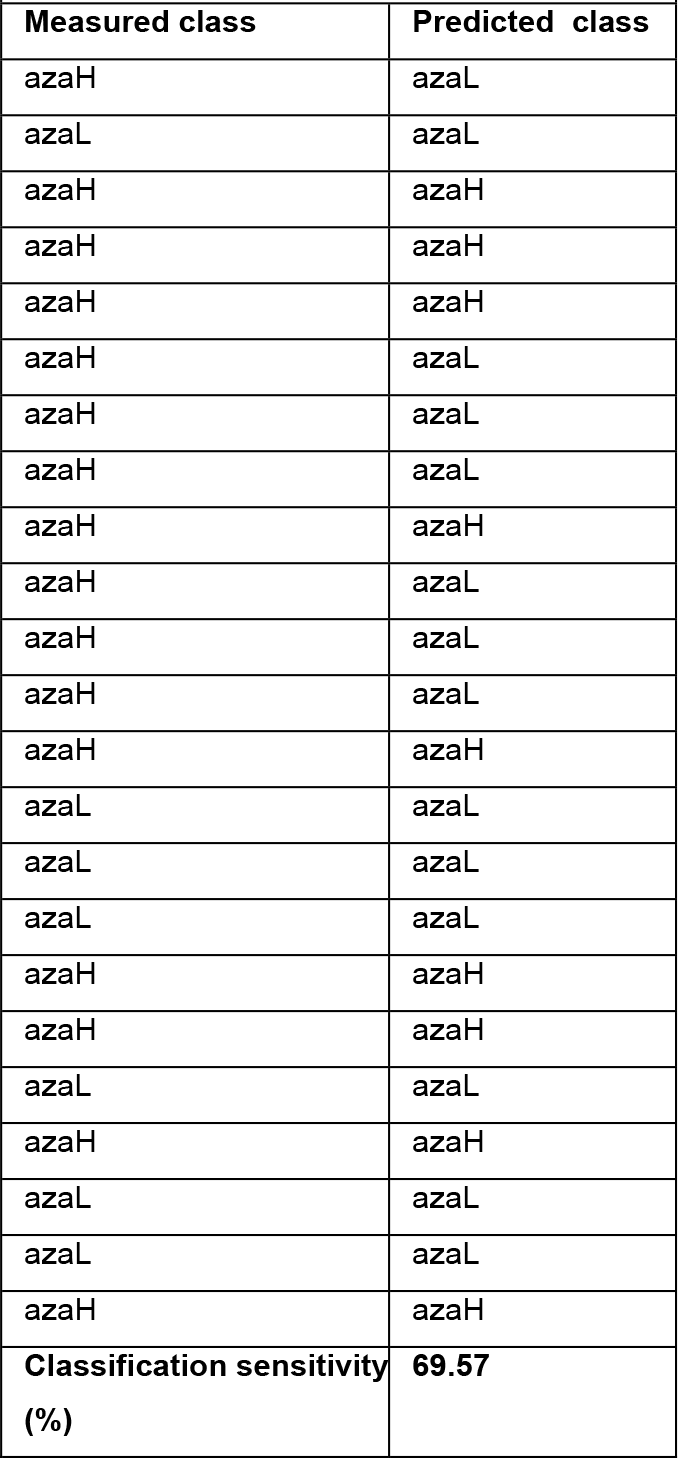
Independent validation using fruit images sampled from an area covering nearly 130,000 sq kms.

## Discussion

### Significance of deep learning approaches in biology

Deep learning methods are producing impressive results across several domains, especially, visual and auditory recognition [16]. Data-intensive biological problems are well suited to be solved using deep learning methods [17]. Biologically inspired neural networks are a class of machine learning algorithms that enable learning from data. Deep learning requires a neural network with multiple layers. Based on the nature of data and the type of question being asked, deep learning methods use either supervised, or unsupervised, or reinforcement learning-based training models. CNNs or ConvNets are multi-layered neural networks trained with back-propagation algorithm, used in recognizing images with minimum pre-processing. In 1998, LeCun and co-workers had described a machine learning technique that was built to learn automatically with less hand-designed heuristics [18]. This formed the basis for development of the CNN field. CNNs combine three architectural ideas: local receptive fields, shared weights and, sometimes, spatial or temporal sub-sampling [19]. Biological data is often complex, multi-dimensional and heterogeneous. However, the possibility of using deep learning methods to discover patterns in such large and complex biological datasets is promising. Till date, image-based neural networks have been used to find patterns that aid disease detection and diagnosis [20–22], secondary structure prediction in proteins [23–25], effects of single nucleotide variation in genes [26], plant [27] and plant stress, phenotyping [28–29], and other biological applications.

### Metabolite concentration class as a test case for deep learning

Plant-based metabolites offer a cheaper, non-toxic and environment-friendly alternative to chemical pesticides. Azadirachtin, derived from the fruits of neem tree, is one such metabolite with known anti-feedant and insect-repellent properties [5]. Seed kernels derived from the tree’s fruits are the primary source of commercial azadirachtin. Neem seeds are collected from naturally grown trees and although inexpensive, the current process of extraction is not scalable due to their limited seed availability in the market. Additionally, azadirachtin content varies a lot depending on the geography of the tree locations, quality of the seeds, age of the trees, soil conditions and climatic factors [30–34], thus causing the commercially available seeds to yield varying levels of azadirachtin. A previous study has reported high variation, as high as 6-fold, in azadirachtin-A and azadirachtin-B content across seed kernels [31]. To circumvent this, total organic synthesis of azadirachtin was attempted in the past [35]. However, the process was time-consuming and produced with a low yield [36]. Recent successful advances in synthetic biology such as producing isoprenoids like beta-Farnesene and the antimalarial drug, artemisinin, in heterologous hosts [1, 37–39] have resurrected the hope for bulk production of metabolites, independent of their natural source. However, synthetic biology requires full knowledge of the metabolite synthesis pathway, and optimization of the pathway and production process within the heterologous host(s). Future work on the elucidation of metabolite production pathways based on genome and transcriptome sequences of neem tree [40–42] coupled with prior knowledge on synthetic biology of terpenoid biosynthesis pathways will pave ways to produce cheaper, more effective and potent azadirachtin. Until then, identifying high azadirachtin-bearing fruits and selecting them as commercial sources will greatly boost the production of high-grade azadirachtin. In the current study, we developed a mobile application that can help collectors in the field to identify trees with high azadirachtin content class quickly, and without knowledge of any laboratory procedures. Although simple, this can have a profound impact on the overall efficiency of the industry.

### Advantages of a predictive approach to identifying metabolite content class

Current methods for metabolite screening, although precise, are cumbersome and require a laboratory equipped with expensive equipment, reagents and expert personnel. Additionally, these methods require all collected fruits to be transported to a central laboratory, thus making the process lengthy and costly. Often, a method that provides a less precise readout of the azadirachtin level, for e.g. low or high, is sufficient to make an initial choice before commercial extraction. The current study is a step towards this aim, and provides a simple, inexpensive and on-the-field method that can be used by anyone, even with little or no training at the collection sites to selectively choose high azadirachtin yielding fruits.

India has nearly 337 million smartphone users, which is expected to grow to 490 million by 2022 [43]. The country is also one of the fastest growing markets for mobile device-based delivery platforms for services and e-commerce. Therefore, we explored the possibility of devising a mobile application can be used at the collection site to determine whether a plant organ has high or low metabolite content. The potential of such an application based on deep learning technology to classify images and predict biological measurements is very enticing. In the current report, we show that it is possible to use an image of a fruit or a leaf to determine whether the intracellular concentration of an analyte within the corresponding cells is high or low. Our method provided us with nearly 70% validation with the fruit model. Since the seed kernels from fruits are the primary source of azadirachtin, our model is well suited to pre-select fruits that can be used as commercial sources. Exciting possibilities similar to the concept demonstrated here can be explored. For example, one can use images to bring quality (and desired trait) control in the fruit and vegetable industry. There are other requirements in the horticultural and agricultural industry that can also be met by the use of image-based learning to deduce characteristics of a certain trait.

### Importance of using both the fruit and the leaf models for classification

As shown in the results section (Figure 4, Table 5), azadirachtin content class predictions using both the fruit and the leaf models together, rather than using any one, yielded fewer false positives, none of which belonged to the azaH class. However, doing so also resulted in lower true positives, although the effect was marginal. The possibility of zero false positives, even at a cost of discovering fewer azaH trees is particularly advantageous and relevant for the azadirachtin extraction industry. Therefore, we recommended the use of both models with the AZA classifier app for sampling in the field. However, there is a possibility that with increased training and improvised models in the future, the classification may work optimally with the fruit model alone, without losing sensitivity or specificity of detection.

### Limitations of our study and future directions

The prediction accuracy of our classier app was not 100%, and we got errors of 19.13% and 15.11%, and 8% and 26.67%, each, using the fruit and leaf models for the low and high metabolite class, respectively. We did not observe an increasing trend of misclassifications towards the aza borderline cases, however (*Figure S1*). This was expected because CNNs are non-parametric classifiers and by definition, no statistical parameters are needed to separate the image classes, even though they were formed based on quantitative thresholds [44–45]. Since the learning of images is not a gradual process that becomes less accurate at a certain end of the azaL or azaH quantitative spectrum, and rather is categorical with respect to the aza classes, we believe that the misclassifications might have resulted from the image-related reasons. Although our initial models are promising for image-based metabolite class prediction, there are several limitations of our study. The performance of any machine learning approach is dependent on both, adequacy of the sampled data and the complexity of the underlying model. In cases where the training set data is limiting, chances of the model over fitting to the training data and not generalizing to other ‘unseen’ data are greater, while in cases where the model is over-simplified or over-regularized, chances of the model under-fitting the data are greater. The first limitation of our study is the relatively narrow size of the training set and lack of diversity of conditions in which images were taken, in terms of brightness, camera magnification, blur, left-right flip, top-bottom flip, degree of rotation, etc, Therefore, it is possible that the model might be an over-fit, and is not a suitable predictor for unseen data. Although we have attempted to lower the impact of this limitation by augmenting the training set with synthetic distortions, the prediction accuracy of AZA classifier is expected to increase with naturally augmenting the training set, in both size and diversity, via further sampling, imaging and metabolite content analyses of neem leaves and fruits. In doing so, the aim would first be to utilize the additional data to validate the underlying model parameters, and second, to enhance the training set, in an iterative manner to minimize both, cross-validation and true validation errors. The second limitation of our study is the lack of metabolite data from individual fruits and leaves of a tree. Despite collecting 10 samples per tree (5 fruits and 5 leaves), we had single metabolite readout per plant. Results from methods that can measure metabolite from a small amount of tissue (such as a single fruit or leaf), may prove valuable in the future in tackling within tree variability. With a reasonable assumption that the intra-tree variation of metabolite content across fruits and leaves in a single tree is lower than the inter-tree variability, we used the more robust method of measuring metabolite from fruits pooled from a tree. Additionally, using both fruit and leaf models allowed us to predict aza with high specificity.

Although the current work is based on images to predict a specific intracellular metabolite, azadirachtin, our method has potential to be applicable with broad utility to any cell type, including three-dimensional shapes, images and architecture, towards real-time prediction of intracellular analytes responsible for a characteristic taste, smell or any other feature of fruits, leaves or other parts of a plant.

## Supplementary Tables

**Table S1.** Training and classification sensitivities of fruit and leaf images pooling all images (original post background removal and 7 distortions to the background removed images). MobileNet architecture was used for training and classification. Presence of models with >70% classification sensitivity in both azaL and azaH classes is highlighted for fruits (row numbers 4-164).

**Table S2.** Training and classification sensitivities of all fruit and leaf images (original post background removal and 7 distortions to the background removed images). Inception V3 architecture was used for training and classification. Presence of fruit models with >70% classification sensitivity in both azaL and azaH classes are highlighted.

**Table S3.** Training of all original leaf images post background removal and 3 distortions (Crop – 30%, Scale – 30% and Flip – left and right), using the MobileNet architecture. Classification sensitivities are reported for each distortion category. Presence of leaf models with >70% classification sensitivity in both azaL and azaH classes are highlighted.

**Table S4.** Training and classification sensitivities forthefruitandleafimages, individually postbackgroundremovalandpost all the 7 distortions got applied to the background-removed images, using the MobileNet architecture. Presence of models with >70% classification sensitivity in both azaL and azaH classes are highlighted in orange for fruits (row numbers 5-164), and in green for leaves (row numbers 165-324).

## Supplementary Figure

**Figure S1.** Misclassification trend versus measured azadirachtin levels.

## Availability of source code and installation requirements

- Project home page: https://github.com/binaypanda/AZA-classifier/tree/master/v1.1
- Project name: AZA classifier
- Project version: 1.1
- For detailed instruction on installation and other prerequisites, please see the README file on github (https://github.com/binaypanda/AZA-classifier)
- License: GNU GPL
- Requirements for the desktop app: -

- Operating system(s): Tested on Linux v18.04 LTS, Windows – Windows 10 and Mac – macOS Mojave, 10.14)
- Programming languages: Java, Python
- Installation pre-requisites (as per tested environments): Xcode Command Line Tools and Homebrew (for Mac), ~Java 8, Python3.6.7 (for Linux & PC), Python 3.7.2 (for Mac), Python pip 19.0.2, Python modules: TensorFlow 1.12 & Numpy 1.16.1 (for Mac, PC and Linux)
- Requirements for the mobile app:

- Smart phone with Android (v5.1)

## Availability of supporting data and materials

The image data used in this study is available at Figshare (https://figshare.com/s/b07ae3faa5accea7c8c3).

## Declarations

NMK and BP are inventors of a filed patent application (E-2/3560/2018/CHE, Application No 201841019772) based on the work described in the manuscript.

## Author’s Contributions

NMK built the azadirachtin classification models using TensorFlow APIs, conceived and implemented using them via desktop and mobile Android apps to predict azadirachtin class for incoming leaf and fruit images, and built the desktop and mobile apps. BP conceived the idea that leaf and fruit images can classify azadirachtin content class. NMK and BP analyzed the data and wrote the manuscript.

## Acknowledgements

We thank Mohanraj I for help in making the apps with the initial versions of the models, and EID Parry for help in sample collection and azadirachtin quantification.

## References

1. Keasling JD. Synthetic biology: A global approach. Nature 2014;510(7504):218.

2. Minervini M, Abdelsamea MM, Tsaftaris SA. Image-based plant phenotyping with incremental learning and active contours. Ecological Informatics 2014;23:35–48.

3. Rostam HM, Reynolds PM, Alexander MR, Gadegaard N, Ghaemmaghami AM. Image based machine learning for identification of macrophage subsets. Scienti_c reports 2017;7(1):3521.

4. Kovaleva L, Zakharova E, Minkina YV. Phytohormonal Control of Male Gametophyte Growth in the Pollen–Pistil System. In: Doklady Biochemistry and Biophysics, vol. 385 Springer; 2002. p. 193–195.

5. Schmutterer H, Sharma S. The Neem Tree Azadirachta indica A. Juss. and Other Meliaceous Plants: Sources of Unique Natural Products for Integrated Pest Management, Medicine, Industry and Other Purposes. Nematologica 1997;43(1):121–121.

6. Mohanty SP, Hughes DP, Salathé M. Using deep learning for image-based plant disease detection. Frontiers in plant science. 2016 Sep 22;7:1419.

7. Bloice MD, Stocker C, Holzinger A. Augmentor: an image augmentation library for machine learning. arXiv preprint arXiv:170804680 2017.

8. Horstmann CS, Cornell G. Core Java: Advanced Features, vol. 2. Pearson Education; 2013.

9. Abadi M, Barham P, Chen J, Chen Z, Davis A, Dean J, et al. Tensorow: a system for large-scale machine learning. In: OSDI, vol. 16; 2016. p. 265–283.

10. Schmidhuber J. Deep learning in neural networks: An overview. Neural networks 2015;61:85–117.

11. Howard AG, Zhu M, Chen B, Kalenichenko D, Wang W, Weyand T, et al. Mobilenets: Ecient convolutional neural networks for mobile vision applications. arXiv preprint arXiv:170404861 2017.

12. Szegedy C, Liu W, Jia Y, Sermanet P, Reed S, Anguelov D, et al. Going Deeper with Convolutions. In: Computer Vision and Pattern Recognition (CVPR); 2015. http://arxiv.org/abs/1409.4842.

13. Ioe S, Szegedy C. Batch Normalization: Accelerating Deep Network Training by Reducing Internal Covariate Shift; 2015. p. 448–456. http://jmlr.org/proceedings/papers/v37/ioffe15.pdf.

14. Szegedy C, Vanhoucke V, Io_e S, Shlens J, Wojna Z. Rethinking the inception architecture for computer vision. In: Proceedings of the IEEE conference on computer vision and pattern recognition; 2016. p. 2818–2826. http://arxiv.org/abs/1512.00567.

15. Smyth N. Android Studio 3.0 Development Essentials- Android 8 Edition. Payload Media, Inc.; 2017.

16. LeCun Y, Bengio Y, Hinton G. Deep learning. Nature 2015;521(7553):436.

17. Ching T, Himmelstein DS, Beaulieu-Jones BK, Kalinin AA, Do BT, Way GP, et al. Opportunities and obstacles for deep learning in biology and medicine. Journal of The Royal Society Interface 2018;15(141):20170387.

18. LeCun Y, Bottou L, Bengio Y, Ha_ner P. Gradient-based learning applied to document recognition. Proceedings of the IEEE 1998;86(11):2278–2324.

19. LeCun Y, Bengio Y, et al. Convolutional networks for images, speech, and time series. The handbook of brain theory and neural networks 1995;3361(10):1995.

20. Esteva A, Kuprel B, Novoa RA, Ko J, Swetter SM, Blau HM, et al. Dermatologist-level classi_cation of skin cancer with deep neural networks. Nature 2017;542(7639):115.

21. Gulshan V, Peng L, Coram M, Stumpe MC, Wu D, Narayanaswamy A, et al. Development and validation of a deep learning algorithm for detection of diabetic retinopathy in retinal fundus photographs. Jama 2016;316(22):2402–2410.

22. Lakhani P, Sundaram B. Deep learning at chest radiography: automated classi_cation of pulmonary tuberculosis by using convolutional neural networks. Radiology 2017;284(2):574–582.

23. Wang S, Sun S, Li Z, Zhang R, Xu J. Accurate de novo prediction of protein contact map by ultra-deep learning model. PLoS computational biology 2017;13(1):e1005324.

24. Spencer M, Eickholt J, Cheng J. A deep learning network approach to ab initio protein secondary structure prediction. IEEE/ACM transactions on computational biology and bioinformatics (TCBB) 2015;12(1):103–112.

25. Wang S, Peng J, Ma J, Xu J. Protein secondary structure prediction using deep convolutional neural _elds. Scientic reports 2016;6:18962.

26. Quang D, Chen Y, Xie X. DANN: a deep learning approach for annotating the pathogenicity of genetic variants. Bioinformatics 2014;31(5):761–763.

27. Pound MP, Atkinson JA, Townsend AJ, Wilson MH, Grifiths M, Jackson AS, et al. Deep machine learning provides state-of-the-art performance in image-based plant phenotyping. GigaScience 2017;.

28. Pound MP, Atkinson JA, Wells DM, Pridmore TP, French AP. Deep learning for multi-task plant phenotyping. InProceedings of the IEEE International Conference on Computer Vision 2017 (pp. 2055–2063).

29. Ghosal S, Blystone D, Singh AK, Ganapathysubramanian framework for plant stress phenotyping. Proceedings of the National Academy of Sciences 2018;115(18):4613–4618. https://www.pnas.org/content/115/18/4613.

30. Chary P. A comprehensive study on characterization of elite Neem chemotypes through myco_oral, tissuecultural, ecomorphological and molecular analyses using azadirachtin-A as a biomarker. Physiology and Molecular Biology of Plants 2011;17(1):49–64.

31. Kaushik N, Singh BG, Tomar U, Naik S, Vir S, Bisla S, et al. Regional and habitat variability in azadirachtin content of Indian neem (Azadirachta indica A. Jusieu). Current Science 2007;p. 1400–1406.

32. Sidhu O, Kumar V, Behl HM. Variability in neem (Azadirachta indica) with respect to azadirachtin content. Journal of Agricultural and Food Chemistry 2003;51(4):910–915.

33. Singh A, Negi M, Rajagopal J, Bhatia S, Tomar U, Srivastava P, et al. Assessment of genetic diversity in Azadirachta indica using AFLP markers. Theoretical and Applied Genetics 1999;99(1-2):272–279.

34. Priyanka S, Rekha HR, Mithilesh S, Rakhi C. Assessment of age and morphometric parameters of seeds on azadirachtin production in neem seed kernels collected from various ecotypes. RESEARCH JOURNAL OF CHEMISTRY AND ENVIRONMENT 2010;14(1):24–28.

35. Veitch GE, Beckmann E, Burke BJ, Boyer A, Maslen SL, Ley SV. Synthesis of azadirachtin: a long but successful journey. Angewandte Chemie International Edition 2007;46(40):7629–7632.

36. Veitch GE, Boyer A, Ley S. The Azadirachtin Story. Angewandte Chemie (International ed in English) 2008 02;47:9402–29.

37. Meadows AL, Hawkins KM, Tsegaye Y, Antipov E, Kim Y, Raetz L, et al. Rewriting yeast central carbon metabolism for industrial isoprenoid production. Nature 2016;537(7622):694–697. https://doi.org/10.1038/nature19769.

38. Paddon CJ, Westfall P, Pitera DJ, Benjamin K, Fisher K, McPhee DL, et al. High-level semi-synthetic production of the potent antimalarial artemisinin. Nature 2013;496:528–532.

39. Ro DK, Paradise E, Ouellet M, Fisher K, Newman K, Ndungu J, et al. Production of the antimalarial drug precursor artemisinic acid in engineered yeast. Nature -London- 2006 4;440(7086).

40. Krishnan NM, Pattnaik S, Deepak SA, Hariharan AK, Gaur P, Chaudhary R, et al. *De novo* sequencing and assembly of *Azadirachta indica* fruit transcriptome. Current Science 2011 101(12): 1553–1561.

41. Krishnan NM, Pattnaik S, Jain P, Gaur P, Choudhary R, Vaidyanathan S, et al. A draft of the genome and four transcriptomes of a medicinal and pesticidal angiosperm *Azadirachta indica*. Bmc Genomics. 2012 Dec;13(1):464.

42. Krishnan NM, Jain P, Gupta S, Hariharan AK, Panda B. An Improved Genome Assembly of *Azadirachta indica* A. Juss. G3: Genes, Genomes, Genetics. 2016 Jul 1;6(7):1835–40.

43. eMarketer Editors, More than a Quarter of India’s Population Will Be Smartphone Users This Year; 2018. [Online; 3-May-2008]. https://www.emarketer.com/content/more-than-a-quarter-of-india-s-population-will-be-smartphone-users-this-year.

44. Paola JD, Schowengerdt RA. A detailed comparison of backpropagation neural network and maximum-likelihood classifiers for urban land use classification. IEEE Transactions on Geoscience and remote sensing. 1995 Jul;33(4):981–96.

45. Foody GM. Status of land cover classification accuracy assessment. Remote sensing of environment. 2002 Apr 1;80(1):185–201.

