## Supplementary figures and images for "Prediction of a plant intracellular metabolite content class using image-based deep learning"

### Supplementary Figure

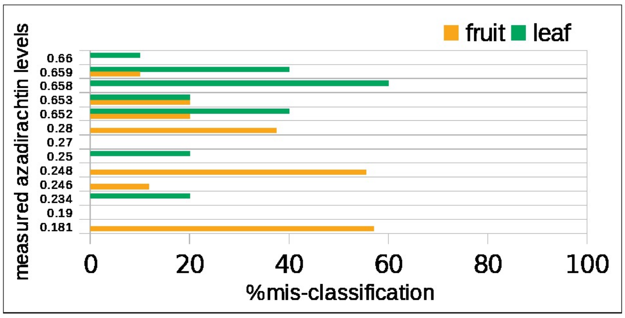
